# Two new species of *Mylochromis* species (Cichlidae) from Lake Malawi, Africa

**DOI:** 10.1101/2024.07.23.604847

**Authors:** George F. Turner

## Abstract

Two new species of *Mylochromis* Regan 1920 are described from specimens collected on shallow rocky habitats on the northwestern coast of Lake Malawi. The generic designation is based on their prominent oblique striped pattern and lack of any of the unique features of other Malawi cichlid genera with this pattern. *Mylochromis rotundus* sp. nov. is distinguished from most congeneric species by its relatively deep, rounded body and lack of enlarged pharyngeal teeth. It is further distinguished from *M. semipalatus* and *M. melanonotus* (if they are different species) by its relatively shorter snout. *Mylochromis durophagus* sp. nov. is distinguished from most congeneric species by its strongly molarised pharyngeal dentition. It differs from *Mylochromis mola* in having a shorter snout, less acutely pointed head profile, deeper body and in having a more continuous oblique stripe. *Mylochromis durophagus* has a much less steep head profile than *Mylochromis anaphyrmus* or *Mylochromis sphaerodon*. It is suggested that *Mylochromis rotundus* was previously identified informally as *Mylochromis sp*. ‘mollis north’, and *M. durophagus* as *M. sp*. ‘mollis chitande’. Lectotypes are designated for comparator species *Mylochromis sphaerodon* and *Mylochromis subocularis*.

## 1 INTRODUCTION

Lake Malawi’s cichlid fish fauna presents a challenge to researchers (Snoeks 2004) as a result of the exceptional number of species (around 800: Konings 2016), many of which are extremely similar morphologically (Turner et al. 2001). The rate of species description is slow and many species, even some well-known ones, remain undescribed, rendering them ineligible to receive IUCN redlisting, or incorporation into standard reference systems such as FishBase, GBIF etc.

Most Malawian cichlids are haplochromines, closely related to the riverine *Astatotilapia calliptera* (Günther, 1894) (Eccles & Trewavas 1989; Joyce et al. 2011; Malinsky et al. 2018). Among Lake Malawi haplochromines, a number of species show an oblique stripe along the flanks, from the nape to the caudal peduncle or caudal fin; this pattern is believed to be unique among haplochromine cichlids (Eccles & Trewavas 1989). In the last major revision of a large section of the Malawian haplochromines, Eccles & Trewavas (1989) assigned 22 of the oblique-striped species to 10 genera *Aristochromis* Trewavas, 1935, *Buccochromis* Eccles & Trewavas, 1989, *Caprichromis* Eccles & Trewavas, 1989, *Champsochromis* Boulenger, 1915, *Corematodus* Boulenger, 1897, *Docimodus* Boulenger, 1897, *Lichnochromis* Trewavas, 1935, *Platygnathochromis* Eccles & Trewavas, 1989, *Taeniolethrinops* Eccles & Trewavas, 1989 and *Tramitichromis* Eccles & Trewavas, 1989, largely on the basis of specialised head and jaw morphology-in some cases these genera also included species with different melanin patterns (*Corematodus, Tramitichromis*). The remaining 16 species were placed in the genus *Maravichromis* Eccles & Trewavas 1989, type *Haplochromis ericotaenia* Regan, 1922, but they were unable to identify any synapomorphic characters for the genus, and its definition was based on the shared possession of the oblique-striped pattern and the lack of putatively apomorphic traits used to define the other genera incorporating oblique-striped species. *Maravichromis* was later found to be a subjective junior synonym of *Mylochromis* Regan, 1920 (Derijst & Snoeks, 1992). The genus was originated in a footnote to Regan’s paper on Tanganyikan cichlids, with the type species *Chromis lateristriga* Günther, 1864, but in his paper on Malawian cichlids published in 1922, that species was included in the genus *Haplochromis* Hilgendorf, 1888, and *Mylochromis* was not listed in the synonymy. No subsequent revision of *Mylochromis* has taken place, and the definition for *Maravichromis* has been assumed to be rolled over to *Mylochromis*.

Subsequently, however, Konings (1993a) revised the genus *Sciaenochromis* Eccles & Trewavas 1989, expelling two species to *Mylochromis*, then proposing that *Mylochromis semipalatus* (Trewavas, 1935) should be considered a junior synonym of *Platygnathochromis melanonotus* (Regan, 1922), that *Platygnathochromis* in turn is a junior synonym of *Mylochromis* (Konings, 1993b) and also that *Chromis subocularis* Günther, 1894 be moved from *Placidochromis* to *Mylochromis* (Konings 1995). Turner & Howarth (2001) described a further two species. However, the synonymy of *M. semipalatus* with *M. melanonotus* was rejected by Snoeks & Hanssens (2004), although maintained by Konings (2016). Fricke et al. (2024) currently accept the validity of *M. semipalatus*, bringing the current total number of species to 22. Snoeks and Hanssens (2004) suggested the genus was in desperate need of revision, that additional undescribed species were intermediate between this genus and *Sciaenochromis* and *Stigmatochromis* Eccles & Trewavas 1989 and that in addition to the three species then recognised to have heavily molariform pharyngeal dentition, they tentatively identified an additional 11 morphological groups. Many probably undescribed species are illustrated in underwater photographs by Konings (2016), but in most cases there are no specimens available to examine. Clearly, much further work is required.

During the examination of a collection of material supporting a programme of genome sequencing (see Malinsky et al. 2018; Svardal et al 2019; Turner et al. 2022 for preliminary results), two new species were identified which fall within the current definition of the genus *Mylochromis* and are formally described in the present work.

## 2 METHODS

Specimens of the new species examined were in existing museum collections at Cambridge University. Counts and measurements were carried out following the methods of Snoeks (2004), using digital callipers and a low power magnifying desk lamp, with additional examination using a binocular dissecting microscope. Angles were measured from photographs using an online protractor (ginifab.com/feeds/angle_measurement/).

Additional material: ***Mylochromis anaphyrmus*** (Burgess & Axelrod, 1973) BMNH 2024.6.25.1-4: 4 specimens 121.3-130.7mm SL, trawled from 60m depth at Chilinda, SE Arm Lake Malawi (−13.743, 35.053), coll. G. Turner 1991; ***Mylochromis chekopae*** Turner & Howarth, 2001: BMNH 1996.10.14.98, holotype, male, trawled from 20m depth off Chekopa on the eastern shore, SE Arm, Lake Malawi (13º 53’ S, 35º 07’ E), Turner, 19 November 1991; BMNH 1996.10.14.99-120, paratypes, 22 specimens, 103.0-122.4 mm SL, collection data as holotype; ***Mylochromis ericotaenia*** (Regan, 1922): BMNH 1921.9.6.148-149 (1) lectotype of *Haplochromis ericotaenia*, 57.9mm SL, collected Lake Nyasa by Wood 1920; BMNH 1935.6.14.2412-2414. 4 specimens, 63.4-81.3mm SL, coll. Lake Nyasa, Christy 1925; BMNH 1935.6.14:2381-2385. 6 mature males 116.1-162.7mm SL, coll. Deep Bay (Chilumba), Malawi, Christy 1925; BMNH 1935.6.14.2401-2402, 2 apparent mature males, 131.9-137.9mm SL coll. Deep Bay, Christy 1925; ***Mylochromis labidodon*** (Trewavas, 1935): BMNH 1935.6.14.2417, lectotype of *Haplochromis labidodon*, 100mm SL, collected from Mwaya, Lake Nyasa, Tanzania by Christy 1925; BMNH 1935.6.14.2418-9, paralectotypes: 3 specimens: one dissected, others 84.9, 85.4 (both very soft & faded), collected with lectotype; BMNH 1935.6.14.2416, paralectotype, 1 specimen 149.9mm SL from Deep Bay (Chilumba), Malawi, coll. Christy 1925; ***Mylochromis mola*** (Trewavas, 1935) 3 specimens: BMNH 1935.6.14.2357, lectotype of *Haplochromis mola*, 139.0mm SL, collected Vua, Lake Malawi, Christy 1925; BMNH 1935.6.14.2358-2359, paralectotypes 110.2-122.4mm SL, collected with lectotype; UMZC 2016.38.43 (field ID: D10-J02), 1 specimen 89.2mm SL, collected Chiofu Bay (−13.533, 34.866), SCUBA, 26 Feb 2016; ***Mylochromis mollis*** (Trewavas, 1935) 2 specimens: BMNH 1934.6.14.1334, lectotype of *Haplochromis mollis*, 131.9mm SL, collected Monkey Bay, Christy 1925; BMNH 1934.6.14.1335: paralectotype 85.0mm SL, collected with lectotype; ***Mylochromis obtusus*** (Trewavas, 1935), BMNH 1935.6.14.1453, holotype of *Haplochromis obtusus*, 190mm SL, coll. SE Arm between Bar and Nkhudzi, Christy 1925; ***Mylochromis sphaerodon*** (Regan, 1922) 2 specimens: BMNH 1921.9.6.146, lectotype (here designated) of *Haplochromis sphaerodon*: figured by E. Fasken in Eccles & Trewavas 1989, 90.2mm SL, collected ‘Lake Nyasa’ by Wood; paralectotype 90.9mm SL, collected with lectotype; ***Mylochromis subocularis*** (Günther, 1894): BMNH 1893.11.15.33, lectotype of *Chromis subocularis* (here designated), figured by Günther 1894 (drawing) and Eccles & Trewavas 1989 (photograph): 94.8mm SL, collected Lake Nyasa & Upper Shire River, presented by H.H. Johnston; BMNH 1935.6.14.1180-1189, 19 specimens 82.3-130.4mm SL, coll. Christy, Bar House, Lake Malawi; BMNH 1966.7.25.15. 1 adult male, 113.5mm SL, collected from Liwonde, Upper Shire River below barrage, by Malawi Fisheries Research Unit/E. Trewavas. UMZC 2021.42.7 (field ID: D23-D03) adult male, not measured, collected from Chirimila catch, Thumbi West Island, Malawi, 29 Jan 2017; UMZC uncatalogued (field code: D12-E09), adult male, not measured (photo only), trawled at 20m depth off Makanjila (−13.769, 34.985), 2 March 2016.

## 3 RESULTS

### Mylochromis rotundus new species

Figures 1-3

**FIGURE 1:**
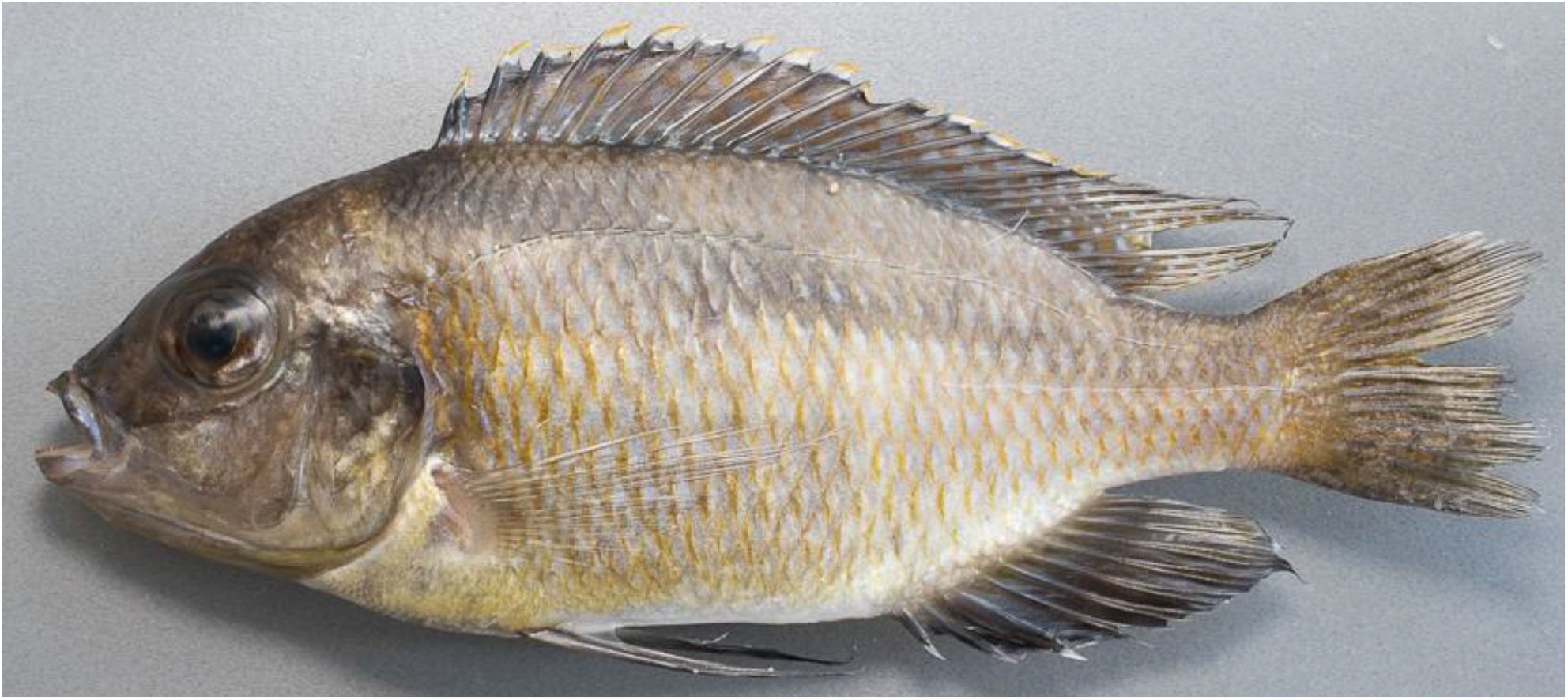
*Mylochromis rotundus*, holotype, UMZC 2016.25.9, 99.7mm SL male.

**FIGURE 2:**
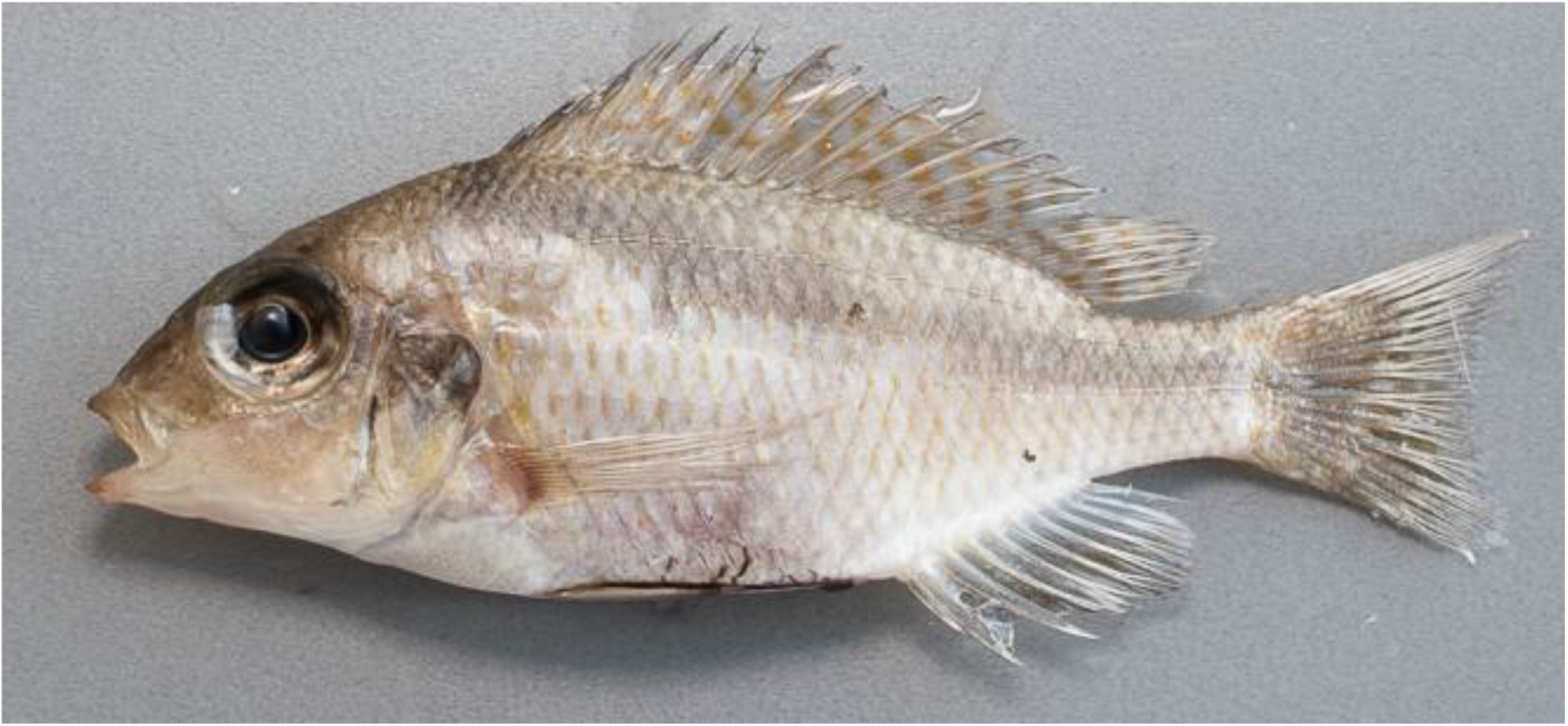
*Mylochromis rotundus*, paratype, UMZC 2016.25.10, 71.5mm SL female.

**FIGURE 3:**
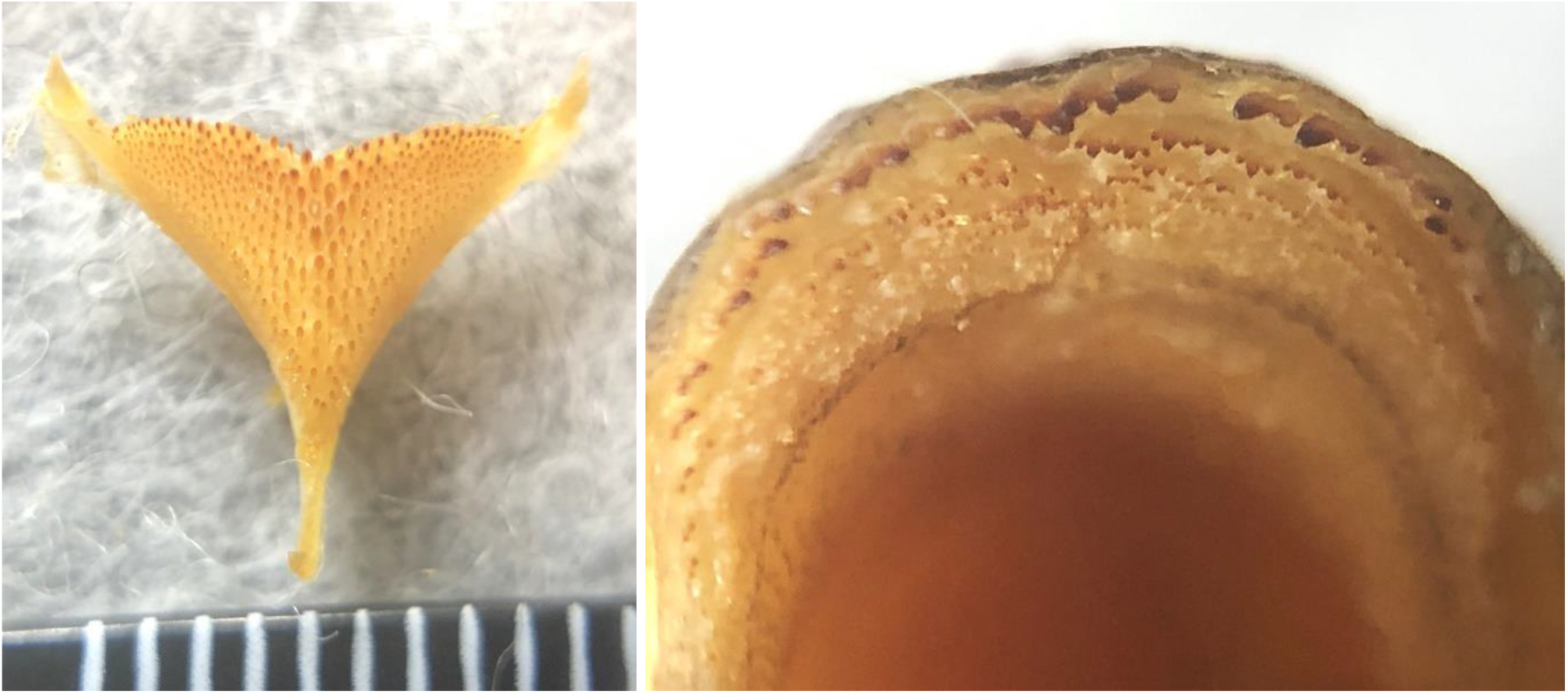
*Mylochromis rotundus*, holotype, lower pharyngeal (left) and oral dentition (lower jaw, right).

urn:lsid:zoobank.org:act:54696C3B-768C-4405-BDA6-9777A3F3A57D

#### Holotype

University Museum of Zoology, Cambridge: UMZC 2016.25.9 apparent male, 99.7mm SL, collected by SCUBA at Mphanga Rocks 23 Feb 2016 by Malawi Cichlid Genomic Diversity Survey (MCGDS).

#### Paratypes (4)

Three apparent males: UMZC 2016.5.4, 88mm SL; 2016.5.5 (lower jaw damaged), UMZC 104.8mm SL; UMZC 2016.5.11, 93.5mm SL and one female UMZC 2016.25.10, 71.5mm SL collected with holotype.

#### Etymology

‘rotundus’ = round, referring to the rounded body shape and short, rounded snout in particular

#### Diagnosis

the lower jaw dentition is ‘*Placidochromis*-type’, with the outer series extending relatively straight towards the posterior end of the jaw. The rounded body, short snout, and large eye enable the species to be readily distinguished from most other known oblique-striped species. Body depth of >38% SL and the lack of enlarged pharyngeal teeth distinguishes the species from all other known congeneric species apart from *M. semipalatus* and *M. melanonotus* (if they are different species). The latter has a distinctively flattened lower jaw at the symphysis. These both have much longer snouts (see Figure 9). In *M. melanonotus*: snout length 1-5 to almost 2x eye diameter (Eccles & Trewavas 1989), *M. semipalatus* snout 1.5x eye diameter (Eccles & Trewavas 1989) v 0.9-1.2x in *M. rotundus*.

#### Description

Body measurements and counts are presented in table 1. *Mylochromis rotundus* is a medium-sized (<105mm SL), laterally-compressed (maximum body depth 2.5-2.6 times maximum width) cichlid fish with a rounded head profile, terminal mouth and large eyes (31.8-38.8% HL) (Figures 1-2). When a melanin pattern is visible, it is dominated by a broad continuous oblique stripe.

**TABLE 1.** Morphometrics and meristics of *Mylochromis rotundus sp. nov*., holotype and 4 paratypes.

|  | Holotype | Paratypes:<br>Mean | Min | Max |
| --- | --- | --- | --- | --- |
| Standard Length (SL, mm) | 99.7 | 89.5 | 71.5 | 104.8 |
| **As % SL** |  |  |  |  |
| Body Depth | 40.7 | 39.8 | 38.6 | 40.4 |
| Head Length | 32.2 | 32.9 | 31.9 | 35.0 |
| Dorsal Fin Base Length | 56.5 | 55.9 | 53.4 | 57.6 |
| Anal Fin Base Length | 19.1 | 18.7 | 18.3 | 19.1 |
| Predorsal Length | 37.2 | 38.1 | 36.8 | 40.4 |
| Preanal Length | 71.2 | 69.9 | 68.7 | 71.8 |
| Prepelvic Length | 34.8 | 34.2 | 32.1 | 36.5 |
| Preventral Length | 42.5 | 40.6 | 38.8 | 42.0 |
| Caudal Peduncle Length | 14.1 | 14.8 | 13.6 | 15.9 |
| Caudal Peduncle Depth | 13.1 | 12.4 | 11.5 | 13.4 |
| **As % Head Length** |  |  |  |  |
| Head Width | 49.2 | 48.6 | 44.4 | 51.0 |
| Interorbital Width | 29.6 | 26.9 | 22.8 | 28.8 |
| Snout Length | 37.7 | 36.0 | 34.8 | 37.4 |
| Lower Jaw Length | 35.2 | 33.8 | 30.6 | 38.0 |
| Premaxillary Pedicel Length | 28.0 | 26.6 | 25.2 | 27.8 |
| Cheek Depth | 23.1 | 17.0 | 16.3 | 17.6 |
| Eye Diameter | 31.8 | 34.8 | 32.0 | 38.8 |
| Lachrymal Depth | 26.2 | 23.9 | 21.2 | 25.3 |
| **Ratio** |  |  |  |  |
| Max. Depth/Head Width | 2.57 | 2.49 | 2.46 | 2.54 |
| Caudal Peduncle Len/Depth | 1.08 | 1.20 | 1.11 | 1.31 |
| **Meristics** |  |  |  |  |
| Upper Gillrakers | 3 |  | 4 | 5 |
| Lower Gillrakers | 13 |  | 10 | 11 |
| Dorsal Fin Spines | 16 |  | 16 | 17 |
| Dorsal Fin Rays | 12 |  | 10 | 11 |
| Anal Fin Rays | 9 |  | 9 | 9 |
| Longitudinal Line Scales | 36 |  | 32 | 33 |
| Cheek Scales | 4 |  | 3 | 3 |

All specimens are relatively deep-bodied and laterally compressed, with deepest part of body generally around 7th dorsal fin spine. Anterior upper lateral profile convex, 40-45^°^ to horizontal anteriorly, slightly concave above eye, continuing straight to tip of snout, with no obvious bulge made by premaxillary pedicel. Jaws isognathous to slightly retrognathous, jaw teeth prominent even when mouth closed. Tip of snout well above level of upper insertion of pectoral fin and about level with the bottom of eye. Lower profile curves gently from lower jaw tip to insertion of pelvic fins, then almost straight from pelvics to first anal spine. Mouth relatively small, gape angled about 32-45^°^ to horizontal and lips relatively thin. Posterior end of maxilla well in front of anterior margin of eye. Eye large, circular and generally below the head profile in lateral view. Lachrymal wider than deep with 5 openings.

Flank scales weakly ctenoid, with cteni becoming reduced dorsally, particularly anteriorly above upper lateral line, where they transition into a cycloid state. Scales on chest relatively large (e.g in type, largest scale 2mm v width across insertion of pelvics, 7.6mm). Gradual transition in size from larger flank scales to smaller chest scales, typical in non-mbuna Malawian endemic haplochromines (Eccles & Trewavas 1989). Caudal fin densely-scaled, over at least proximal ¾.

Cephalic lateral line pores fairly inconspicuous and flank lateral line shows usual cichlid pattern of separate upper and lower portions, with 0-4 pored scales after kink in upper lateral line and 2-3 smaller pored scales after line of flexion of hypurals.

Pectoral fins relatively short, extending beyond vent but not to first anal spine, while pelvics occasionally just reach first anal spine base. Filaments of dorsal and anal fins reaching just past base of caudal fin. Caudal emarginate.

Lower jaw relatively sturdy and broad, with marked mental process. Outer series of teeth in lower jaw stout, erect, prominent, subequally bicuspid with 2 large rounded cusps, generally deeply implanted in fleshy gums (Figure 3). Upper jaw outer series similar, but in some specimens relatively long shafts visible. In both upper and lower jaws, 3-5 irregular inner series of relatively large, erect, stout tricuspid teeth with blunt tips.

Lower pharyngeal bone small, lightly-built, Y-shaped, carrying small, short, slender blunt teeth (Figure 3). None notably enlarged, although central teeth tend to be larger than lateral ones. Gill rakers widely spaced, pointed, finger-like with thick bases.

Preserved colouration of female beige, paler on chest and belly, flanks scales with a brownish spot anteriorly. Strong dark oblique stripe from nape to the caudal fin base, extending into fin rays. Stripe is continuous, but has a ‘stepped’ form of thinner lines joining elongated midlateral and suprapectoral blotches. A large dark blotch on upper half of operculum. Dorsal fin with dull orange spots throughout and caudal, anal and pelvic fins dusky. Mature males darker with golden spots on majority of flank scales. Dorsal, caudal, anal and pelvic fins dark, dorsal and caudal with numerous orange spots. Dorsal fin lappets white with orange tips. It is possible that this is not the full breeding dress and that fully courting males are bright blue (Figure 4).

**FIGURE 4:**
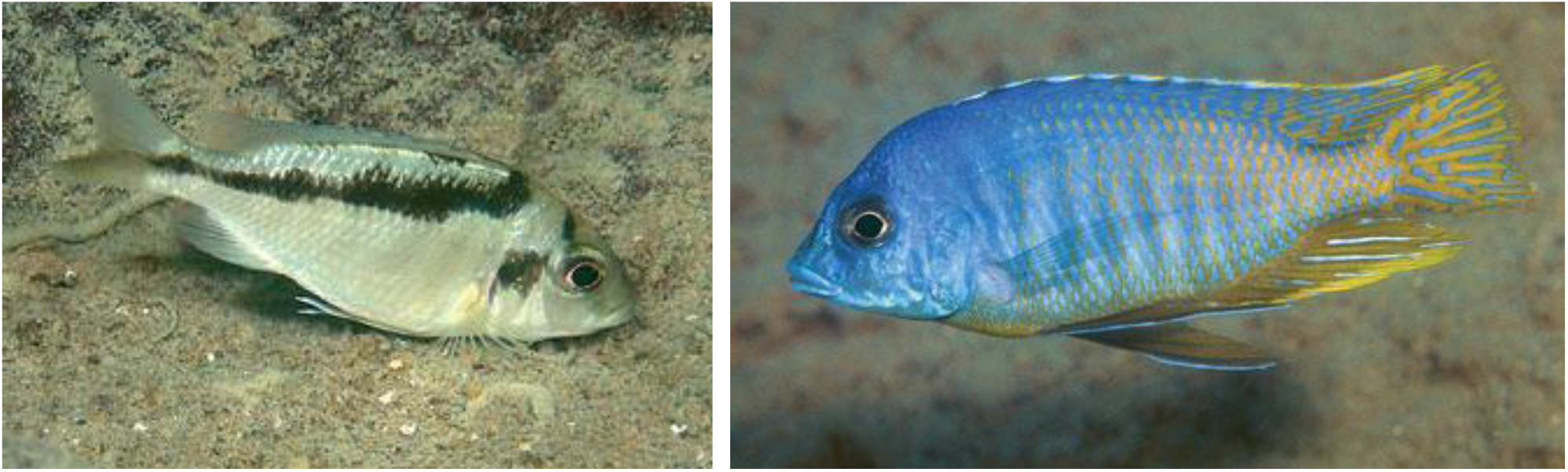
Live coloration of specimens provisionally identified as *Mylochromis sp*. ‘mollis north’ which may represent *M. rotundus*: female/immature from Katale Island, Chilumba (left), mature male from Hora Mhango, NW coast between S. Rukuru River and Ruarwe (right). Photos A. Konings.

#### Distribution and ecology

Known only from a collection from shallow water at Mphanga Rocks near Chilumba in the north-western part of Lake Malawi. Photographs under the name *M. sp*. ‘mollis north’, which may or may not be of this species, are shown from the coast between Ruarwe and Chilumba (Konings 2016; Figure 4). It appears to be a species of rocky habitats.

### Mylochromis durophagus sp. nov

Figures 5-8

**FIGURE 5:**
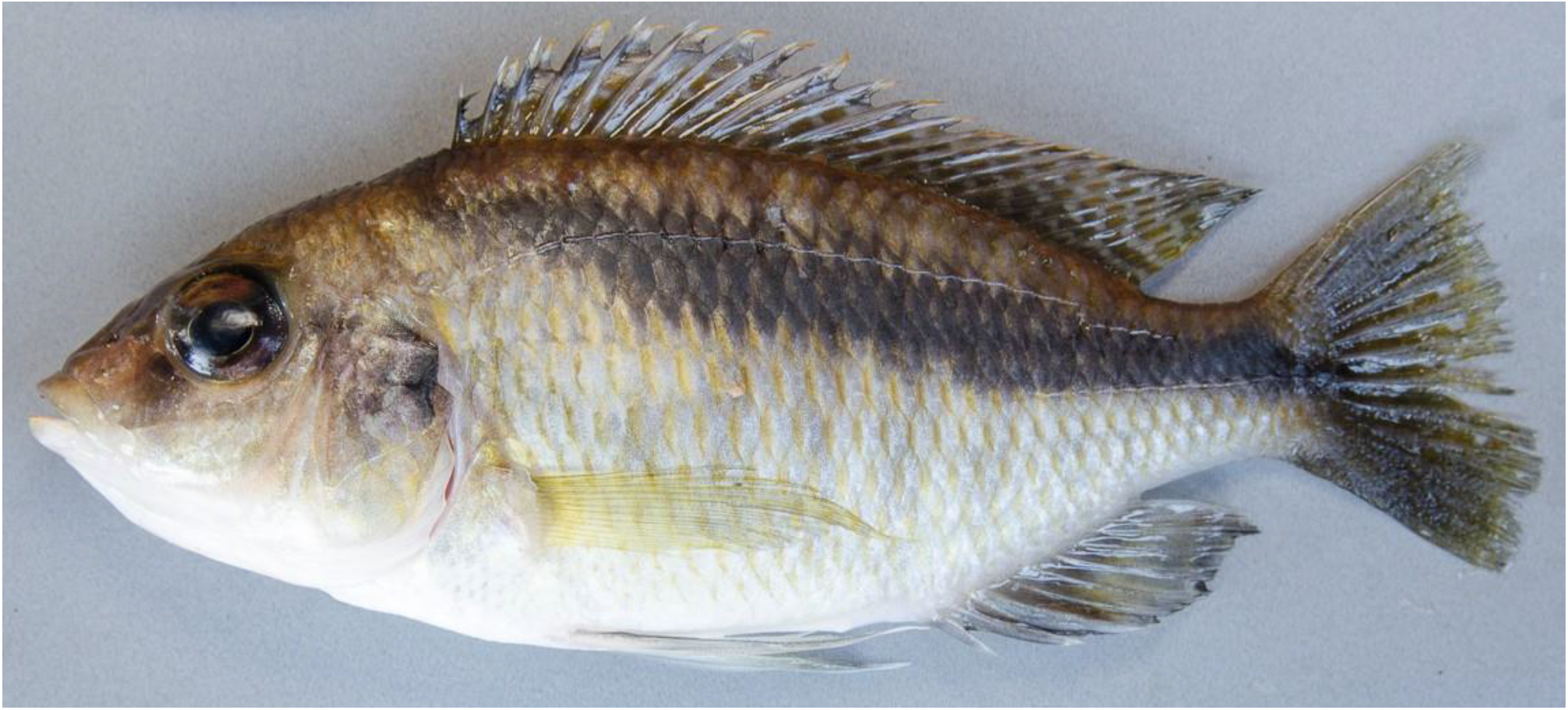
*Mylochromis durophagus*, holotype, UMZC 2016.18.13, 89.7mm SL male, freshly collected from Nkhata Bay 20 Feb 2016 [photo H. Svardal].

urn:lsid:zoobank.org:pub:B29807EB-60A6-4389-A235-993B72E39C9B

#### Holotype

University Museum of Zoology, Cambridge: UMZC 2016.18.13, apparent male, 89.7mm SL, collected by SCUBA at Nkhata Bay 20 Feb 2016 by Malawi Cichlid Genomic Diversity Survey (MCGDS).

#### Paratypes (2)

UMZC 2016.18.15, apparent female, 67.5 mm SL, collected with holotype and one female UMZC 2016.25.2, 80.0mm SL collected by SCUBA at Mphanga Rocks, Chilumba, 23 Feb 2016 by Malawi Cichlid Genomic Diversity Survey (MCGDS).

#### Etymology

‘duro-’ = from durus, Latin, meaning ‘hard’ + ‘-phagus’=, Latin, meaning ‘to eat’, referring to the presumed diet of hard-shelled invertebrates indicated by the molariform pharyngeal dentition.

#### Diagnosis

the lower jaw dentition is ‘*Placidochromis*-type’, with the outer series extending in a relatively straight line towards the posterior end of the jaw. This and the strong oblique stripe and lack of specialised morphology characterises the species as a member of the genus *Mylochromis*. The strongly molarised pharyngeal dentition distinguishes the species from all other known *Mylochromis* except *M. anaphyrmus, M. mola* and *M. sphaerodon* (Figure 9). *Mylochromis mola* differs in having a long snout (36.3-40.3 v 33.4-37.4% HL in *M. durophagus*), more acutely pointed head profile, more slender body (body depth 34.5-37.1 v 39.1-40.2% SL v in *M. durophagus*) and in having a blotchy, interrupted oblique stripe. *Mylochromis durophagus* has a much less steep head profile than *Mylochromis anaphyrmus* (40^°^ v 56-67^°^) and a more upwardly angled gape (40^°^ v 10-20^°^). *Mylochromis durophagus* also has a less steep head profile than *M. sphaerodon* (40^°^ v 50-52^°^).

#### Description

Body measurements and counts are presented in table 2. *Mylochromis durophagus* is a medium-sized (<90mm SL), laterally-compressed (maximum body depth 2.35-2.42 times maximum width) cichlid fish with a rounded head profile, terminal mouth and large eyes (29.3-33.1% HL). Prominent, broad continuous oblique stripe on flanks, narrowing anteriorly (Figures 5-6).

**TABLE 2.** Morphometrics and meristics of *Mylochromis durophagus sp. nov*., holotype and 2 paratypes.

|  | Holotype | Paratypes:' |  |
| --- | --- | --- | --- |
| **Standard Length (mm)** | 89.7 | 67.8 | 80 |
| **As % SL** |  |  |  |
| Body Depth | 40.2 | 39.7 | 39.1 |
| Head Length | 35.8 | 35.7 | 36.9 |
| Dorsal Fin Base Length | 56.6 | 55.2 | 55.3 |
| Anal Fin Base Length | 19.8 | 19.8 | 19.5 |
| Predorsal Length | 37.9 | 39.4 | 39.9 |
| Preanal Length | 71.2 | 70.5 | 71.5 |
| Prepelvic Length | 38.0 | 35.7 | 38.6 |
| Preventral Length | 44.0 | 42.3 | 45.4 |
| Caudal Peduncle Length | 12.6 | 14.6 | 14.4 |
| Caudal Peduncle Depth | 13.5 | 13.0 | 13.3 |
| **As % Head Length** |  |  |  |
| Head Width | 46.7 | 45.9 | 45.1 |
| Interorbital Width | 26.5 | 23.6 | 22.4 |
| Snout Length | 37.4 | 33.5 | 36.6 |
| Lower Jaw Length | 35.2 | 36.4 | 34.2 |
| Premaxillary Pedicel Length | 30.5 | 31.0 | 28.5 |
| Cheek Depth | 16.8 | 16.1 | 18.3 |
| Eye Diameter | 29.3 | 33.1 | 31.9 |
| Lachrymal Depth | 20.9 | 22.7 | 24.1 |
| **Ratios** |  |  |  |
| Max. Depth/Head Width | 2.41 | 2.42 | 2.35 |
| Caudal Peduncle Len/Depth | 0.93 | 1.13 | 1.08 |
| **Meristics** |  |  |  |
| Upper Gillrakers | 5 | 4 | 4 |
| Lower Gillrakers | 10 | 10 | 11 |
| Dorsal Fin Spines | 17 | 17 | 17 |
| Dorsal Fin Rays | 11 | 10 | 10 |
| Anal Fin Rays | 9 | 9 | 9 |
| Longitudinal Line Scales | 33 | 32 | 32 |
| Cheek Scales | 4 | 3 | 2 |

**FIGURE 6:**
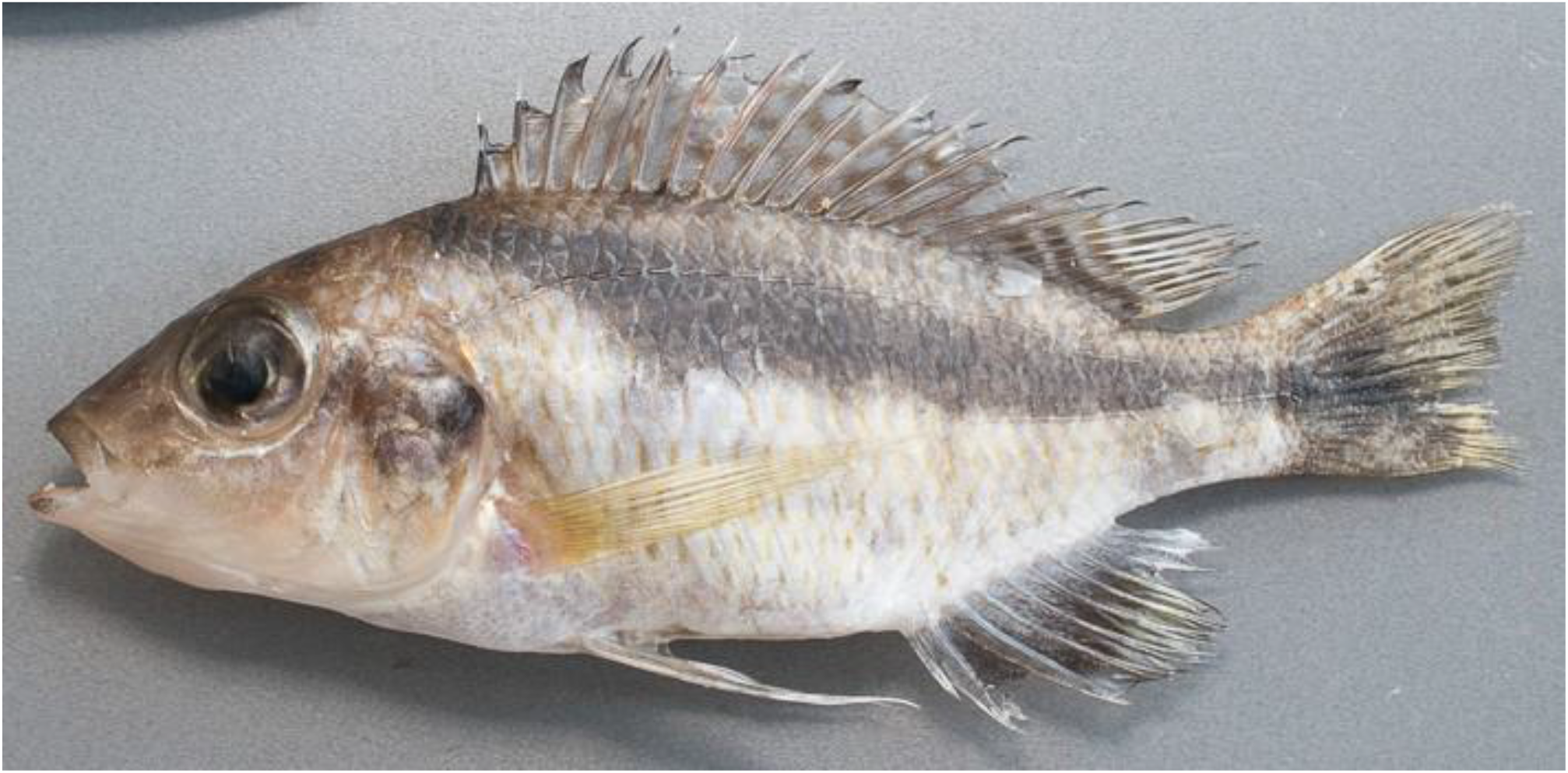
*Mylochromis durophagus*, paratype, UMZC 2016.25.2, 80.0mm SL female, freshly collected by SCUBA at Mphanga Rocks, Chilumba, 23 Feb 2016 [photo H. Svardal].

**FIGURE 7:**
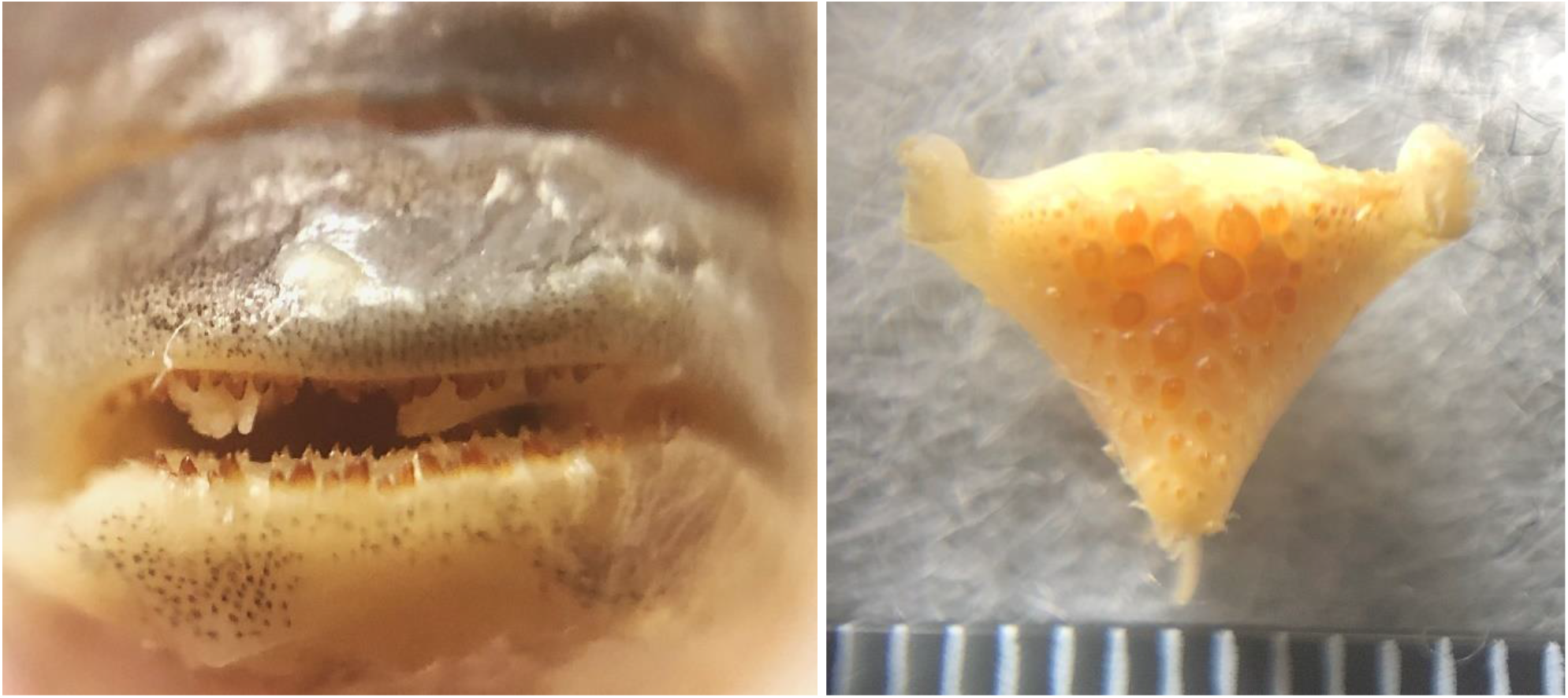
*Mylochromis durophagus*: jaw teeth of holotype (left) and lower pharyngeal bone of 80mm paratype UMZC 2016.25.2.

All specimens are moderately deep-bodied and laterally compressed, with deepest part of body generally around 1^st^-5^th^ dorsal fin spine. Anterior upper lateral profile moderately convex, straight, angled about 40^°^ to horizontal, from tip of snout to above eye, with little or no obvious bulge made by premaxillary pedicel, then curving gently to first dorsal spine. Jaws isognathous to slightly retrognathous, jaw teeth not prominent when mouth closed. Tip of snout well above level of upper insertion of pectoral fin and about level with the bottom of eye or slightly above. Lower profile curves from lower jaw tip to insertion of pelvic fins, then almost straight from pelvics to first anal spine. Mouth relatively small, gape angled about 40^°^ to horizontal. Lips moderately fleshy in larger specimens, relatively thin in smallest. Posterior end of maxilla well in front of anterior margin of eye. Eye large, circular and just below the head profile in lateral view. Lachrymal squarish with 5 openings.

Flank scales weakly ctenoid, with cteni becoming reduced dorsally, particularly anteriorly above upper lateral line, where they transition into a cycloid state. Scales on chest relatively large. Gradual transition in size from larger flank scales, as is typical in non-mbuna Malawian endemic haplochromines (Eccles & Trewavas 1989). Caudal fin densely-scaled, over at least proximal ¾.

Cephalic lateral line pores fairly inconspicuous and flank lateral line shows usual cichlid pattern of separate upper and lower portions, with up to 4 pored scales after kink in upper lateral line and 1-3 smaller pored scales after line of flexion of hypurals.

Pectoral fins relatively long, to around vertical plane through 3^rd^ anal spine, while pelvics occasionally just reach first anal spine base. Filaments of dorsal and anal fins reaching just past base of caudal fin in larger specimens, but well short in smallest. Caudal fin emarginate.

Lower jaw bones relatively slight and lacking a mental process. Outer series of teeth in lower jaw stout, erect, prominent and deeply implanted, bicuspid with pointed, obliquely truncated major cusps. Upper jaw outer series similar, but both cusps more rounded, larger becoming more pointed posteriorly. In both upper and lower jaws, inner teeth very small, scattered, not forming clear rows.

Lower pharyngeal bone heavily built with hemispherical molariform teeth central-posteriorly. Gill rakers widely spaced, thick, blunt.

Female beige, paler on chest and belly, with a strong oblique stripe from nape to the caudal fin base, extending into fin rays. Dorsal fin has dull orange spots in 2 rows on spinous membranes, 3 oblique rows in soft dorsal; lappets dark in spinous portion. Caudal, anal and pelvic fins dusky, pectorals yellowish. Mature male darker with golden spots on majority of flank scales. Dorsal, caudal, anal and pelvic fins dark, dorsal and caudal with numerous orange spots. Dorsal fin lappets white with orange tips. Dark lachrymal stripe visible. Based on other Malawian haplochromines, this is unlikely to be full male breeding dress.

#### Distribution and ecology

Known only from two specimens collected from rocky shores at Nkhata Bay and one at Mphanga Rocks near Chilumba in the north-western part of Lake Malawi. All were collected in shallow water by SCUBA divers.

## 4 DISCUSSION

The two species described above are relatively poorly known and were not adequately distinguished during field collections, but only identified from preserved material examined as part of ongoing genomic studies (for some preliminary results, see Malinsky et al. 2018; Svardal et al 2019; Turner et al. 2022). They were initially recorded as *Mylochromis cf. mollis*. Later investigation of the type material of *M. mollis* showed that this species is more slender in build, while dissection of the pharyngeal bones indicated that we were likely dealing with two species, one with strongly molariform pharyngeal dentition and the other with papilliform dentition. Intraspecific polymorphism in pharyngeal structures is well-known from a Mexican cichlid, *Herichthys minckleyi* (Kornfield & Taylor, 1983) (Bell et al. 2019). Intra-population variation in pharyngeal dentition in African cichlids is relatively poorly known: some variation occurs in the Lake Malawi rocky shore species *Labidochromis caeruleus* Fryer 1956, but the relatively small sample size (Lewis 1982) makes it difficult to know if this is a discrete polymorphism or continuous variation. There is also variation in pharyngeal bones within the Lake Masoko population of *Astatotilapia calliptera*, although much of this is associated with divergence among ecotypes associated with microhabitat preferences (Malinsky et al. 2015; Carruthers et al. 2022). In general, however, it generally appears that such clear differences in lower pharyngeal dentition correspond to species differences rather than polymorphisms (Eccles & Trewavas 1989). Although the sample sizes in the present study are relatively small, the molariform and papilliform individuals also showed non-overlapping differences in Head Length as % SL, Premaxillary Pedicel Length as % Head Length and Maximum Body Depth/ Head Width (Tables 1 & 2), indicating that the two forms represented different morphotypes in unconnected traits consistent with them being different species. As with many other very similar Malawian cichlids, with practice, they could probably be differentiated by a subtle difference in overall appearance, in this case largely from the head profile, without the need for dissection of the pharyngeal bones.

The two new species described above add to the long list of species within *Mylochromi*s. Many undescribed Malawi cichlids have been known via informal names since Ribbink et al. (1983) popularised the practice in their ground-breaking natural history overview of the rocky shore species of the Malawian coast and built on in works such as the successive books by Konings (1989-2016). Konings’ most recent work (2016) lists 38 *Mylochromis* species, of which 17 are apparently undescribed. This is also the number listed on the ‘Cichlid Room Companion’ website (cichlidae.com), which is regularly updated based on both academic and aquarium-hobbyist literature.

It is not always straightforward to match formal descriptions with ‘field species’ shown by Konings and other workers, whose identifications are based largely on underwater photographs. In most cases, details of morphological features such as dentition, gillrakers and pharyngeal jaws are unknown. It is also unclear whether the field identifications of relatively rare species really correspond to the described species. This is attested to by the changing (improving!) names given to same taxon (sometimes the same photograph) across successive edition of Konings’ book. Equally, a number of described species are known from a relatively small number of preserved specimens, and may not all actually represent different species: *M. balteatus* and *M. melanotaenia* are only separated on the key given by Eccles & Trewavas (1989) on the basis of the number of teeth in the outer series of the lower jaw (51-53 v 38-46), although differences in lip thickness and position of lateral stripe are also mentioned in the text. *Mylochromis guentheri* and *M. mollis* appear to differ only in that the former has retrognathous jaws and procumbent lower jaw teeth (Eccles & Trewavas 1989), but this is by no means obvious in all specimens in the type series.

While voucher specimens and formal descriptions give a good insight into morphological traits, live colours may not be captured well. The specimens at the Cambridge Museum were initially collected by the Malawi Cichlid Genomic Diversity Survey (MCGDS) during 2016: a variety of sources and techniques were employed. The material of *M. rotundus* and *M. durophagus* were obtained by divers collecting fish on rocky shores, but these species were not individually observed underwater and were not photographed until post mortem.

Tentatively, it seems that the papilliform *M. rotundus* may be conspecific with Konings’ *M. sp*. ‘mollis north’ and the molarifom *M. durophagus* with Konings’ *M. sp*. ‘mollis chitande’.

*Mylochromis sp*. ‘mollis north’ was recorded from Hora Mhango, Kakusa and Katale Island, all sites north of Ruarwe in the far NW of the lake, while *M. rotundus* was only collected from the Chilumba area, the more southerly sites of Konings were not sampled in the 2016 collecting trip. Konings reports the species as feeding from sand. Males are reported to have a yellow breast and to dig a ‘cave-crater’ bower.

Konings (2016) records *M. sp*. ‘mollis chitande’ from a number of sites around the Chilumba area (Mdoka to Maison Reef), which overlaps with the range of the specimens of *M. durophagus* (Mphanga Rocks, Nkhata Bay). Konings recorded the species as uncommon, and mainly inhabiting the rock-sand interface area (‘intermediate habitat’), picking invertebrates mainly from rocky substrates. He comments on two of the main identification feature of *M. durophagus*, the pointed snout and broad, irregular oblique stripe. Underwater photographs show a similar stripe pattern to *M. durophagus*, in particular, the way the stripe is narrow anteriorly, and then suddenly becomes much wider below around the 6^th^-8^th^ dorsal fin spine. There is also a hint to a large dark blotch on the operculum in the illustrations (figure 8) similar to that seen in the type specimens (figures 5 & 6).

**FIGURE 8:**
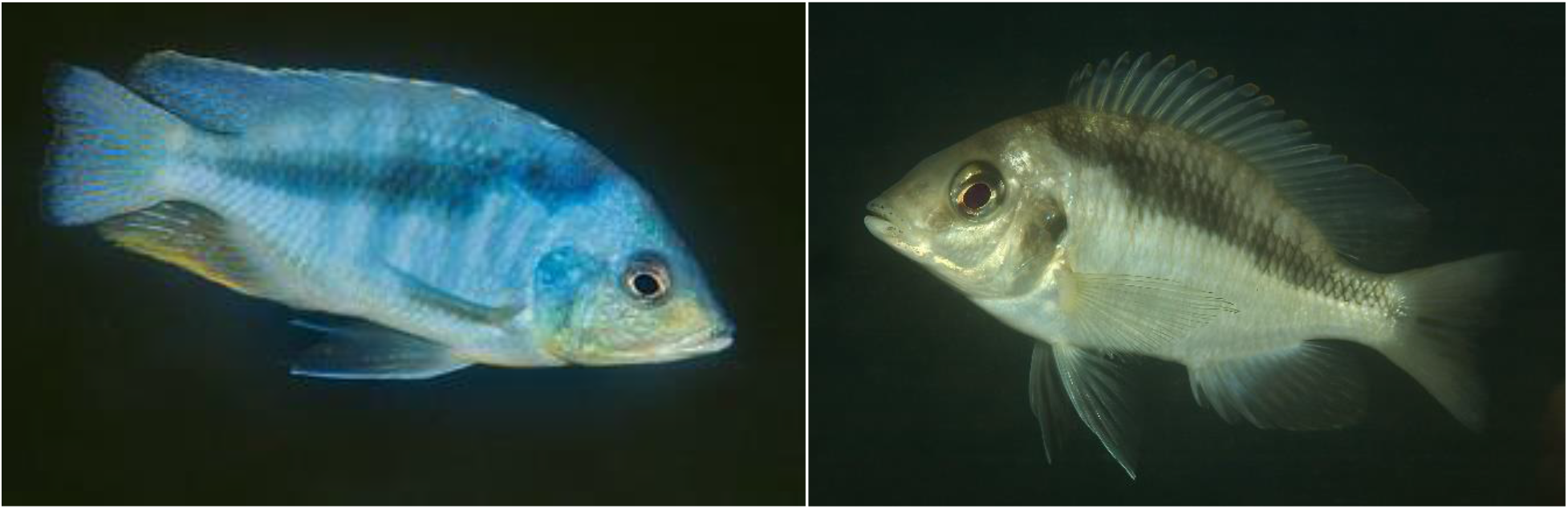
Live coloration of specimens provisionally identified as *Mylochromis sp*. ‘mollis chitande’ which may represent *M. durophagus*: male (left) and apparent female (right) from Maison Reef, Chilumba, NW coast of Lake Malawi (right). Photos A. Konings.

**FIGURE 9:**
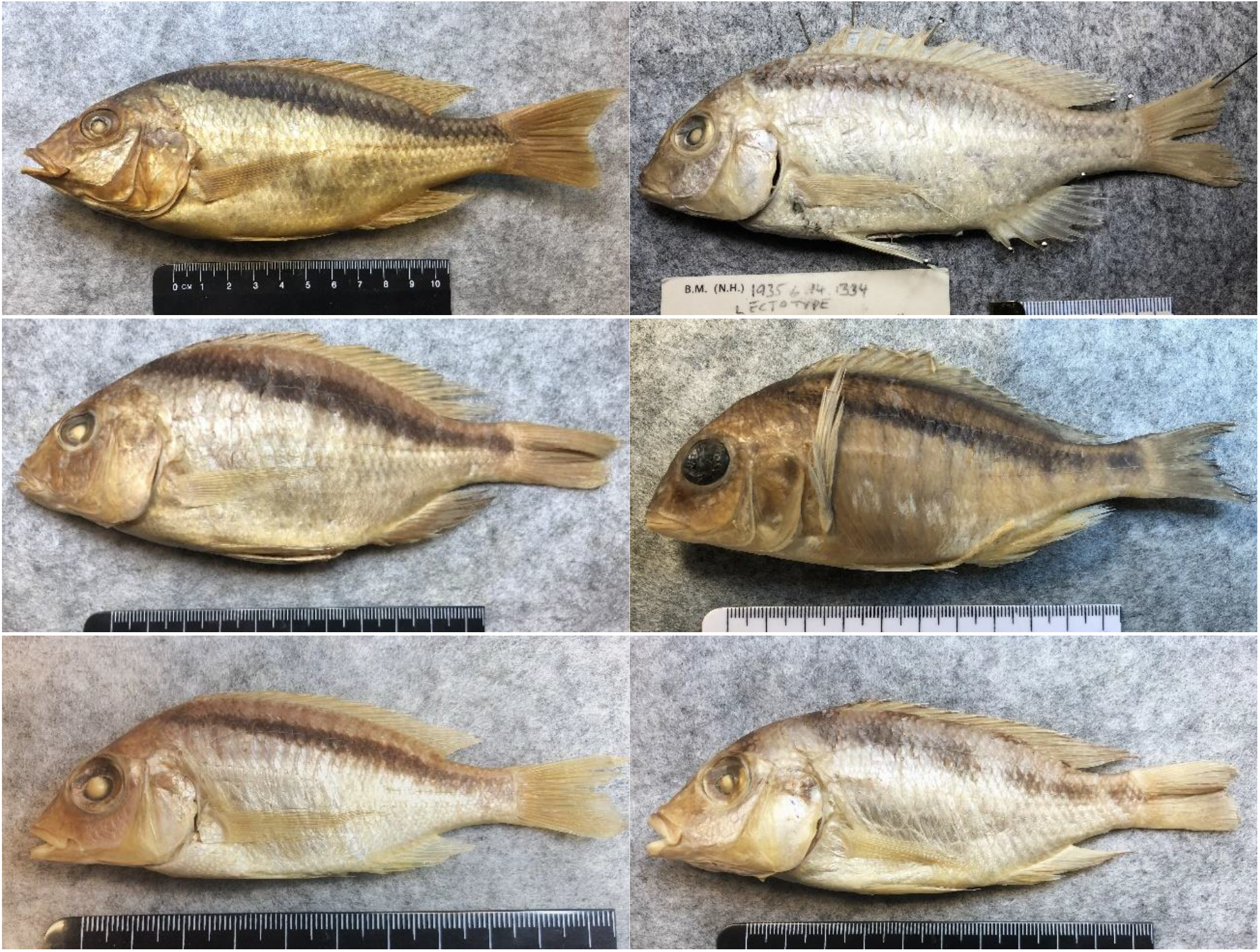
Comparable species: (top left) *Mylochromis melanonotus* lectotype BMNH 1921.9.6.163, 172mm SL; (top right) *Mylochromis mollis* lectotype BMNH 1935.6.14.1334, 131.9mm SL; (centre left) *Mylochromis semipalatus* lectotype BMNH 1935.6.14.1321, 141mm SL; (centre right) *Mylochromis anaphyrmus*, BMNH 2024.6.25.1-4, 130.7mm SL; (bottom left) *Mylochromis sphaerodon* lectotype 1921.9.6.147, 90mm SL; (bottom right) *Mylochromis mola*, BMNH1935.6.14.2357-2359, 122.4mm SL [photos; Turner].

## ACKNOWLEDGEMENTS

I thank Rupert Collins, Oliver Crimmen, James Maclaine and Simon Loader at the Natural History Museum in London for helping with access to specimens and cataloguing new material and Matthew Lowe and Natalie Jones for help with the collection at the Cambridge Zoology Museum. Funding for the 2016 collection was provided by a Wellcome Trust award (WT207492) to Richard Durbin at University of Cambridge and samples exported under an Export Permit from the Department of Fisheries Malawi 20/4/8 and Research Permit from the Government of Malawi NCST/RTT/1/20. Manfred Malinsky, Mexford Malumpwa, Hannes Svardal, Alexandra Tyers helped with the collections at Chilumba and Nkhata Bay in 2016. I am grateful to Hannes Svardal and Ad Konings for permission to use their photographs.

